# Isoflavone treatment attenuates cardiovascular-kidney-metabolic (CKM) syndrome symptoms in an experimental model of ovariectomized rats

**DOI:** 10.64898/2025.12.23.696268

**Authors:** Thais de Souza Lima, Edson de Andrade Pessoa, Amaro Nunes Duarte Neto, Cinthya dos Santos Cirqueira Borges, Márcia Bastos Convento, Bianca Castino, Andréia Silva de Oliveira, Alef Aragão Carneiro dos Santos, Adriana Carbonel, Cassiane Dezoti, Maria de Fatima Vattimo, Fernanda Teixeira Borges

**Affiliations:** Nephrology Division, Department of Medicine, Universidade Federal de São Paulo, São Paulo, SP, Brazil; Interdisciplinary Postgraduate Program in Health Sciences, Universidade Cruzeiro do Sul, São Paulo, SP, Brazil; Experimentation Laboratory in Animal Model, School of Nursing, Universidade de São Paulo, São Paulo, Brazil; Histology and Structural Biology Division, Department of Morphology and Genetics, Universidade Federal de São Paulo, São Paulo, SP, Brazil; Department Medical-Surgical Nursing, School of Nursing, Universidade de São Paulo, São Paulo, Brazil; Department of Pathology, Medical School. Universidade de São Paulo, São Paulo, Brazil; Department of Pathology, Instituto Adolfo Lutz, São Paulo, Brazil

## Abstract

Cardiovascular–kidney–metabolic (CKM) syndrome represents an integrated spectrum of obesity, type 2 diabetes mellitus, cardiovascular disease, and chronic kidney disease, driven by shared metabolic, inflammatory, and fibrotic mechanisms. Menopause-related estrogen deficiency exacerbates these disturbances, increasing susceptibility to metabolic syndrome and renal injury. This study investigated whether dietary supplementation with soy isoflavones attenuates obesity-associated kidney injury in ovariectomized rats exposed to a high-fat, high-caloric diet. Female Wistar rats underwent bilateral ovariectomy and were assigned to control diet, hipercaloric diet (DH), or DH supplemented with isoflavones (DH+ISO). Metabolic, biochemical, oxidative stress, histopathological, and inflammatory parameters were evaluated over 120 days. The DH diet induced weight gain, hypertension, dyslipidemia, hyperglycemia, impaired renal function, increased oxidative stress, and tubulointerstitial injury, consistent with a CKM phenotype. Isoflavone supplementation did not prevent diet-induced weight gain but attenuated systolic blood pressure elevation and improved lipid and glycemic profiles. Renal function was partially preserved in DH+ISO rats, as evidenced by lower plasma creatinine levels, improved creatinine clearance, reduced urea levels, and decreased urinary peroxide excretion compared with DH animals. Histopathological analysis revealed persistent tubular injury in both DH groups, although inflammatory and oxidative markers were attenuated in isoflavone-treated rats. Immunohistochemistry demonstrated reduced renal IL-6 expression in the DH+ISO group, accompanied by trends toward decreased macrophage infiltration and IL-1β expression. Collectively, these findings indicate that soy isoflavones mitigate metabolic, inflammatory, and oxidative pathways implicated in obesity-related kidney injury in a postmenopausal experimental model. Isoflavone supplementation may represent a complementary therapeutic strategy for attenuating CKM-related renal damage associated with modern diets and estrogen deficiency.

## Introduction

Obesity, type 2 diabetes mellitus, cardiovascular disease, and chronic kidney disease constitute an increasingly prevalent and interrelated health challenge. This constellation of conditions has recently been named cardiovascular–kidney–metabolic (CKM) syndrome by the American Heart Association^1^. CKM syndrome is driven by convergent pathophysiological mechanisms, including hyperglycemia, insulin resistance, activation of the renin–angiotensin– aldosterone system, advanced glycation end-product formation, oxidative and endoplasmic reticulum stress, lipotoxicity, mitochondrial dysfunction with impaired energy metabolism and chronic low-grade inflammation.

Obesity arises from a sustained positive energy balance driven by excessive caloric intake and sedentary behavior and is strongly associated with visceral adiposity, systemic inflammation, oxidative stress, and progressive renal dysfunction^2,3^. High-fat diets disrupt energy homeostasis and promote lipid accumulation in metabolically active organs, including the liver and kidney, thereby predisposing to chronic kidney disease (CKD). Experimental models based on high-calorie diets are widely employed to investigate obesity-related kidney injury due to their close resemblance to the metabolic disturbances observed in humans.

Fructose intake, particularly when combined with a high-fat diet, further exacerbates metabolic and renal injury^4^. Excessive fructose consumption, common in sugar-sweetened beverages and processed foods, accelerates the development of obesity and metabolic syndrome (MS)^5,6^. Experimental and clinical evidence indicates that fructose intake induces albuminuria, alters blood pressure regulation, and worsens metabolic parameters, including triglyceride levels and glycated hemoglobin^7^.

At the renal level, fructose exerts early direct effects on proximal tubular cells, stimulating cellular proliferation and metabolic stress. Chronic exposure promotes tubulointerstitial injury and accelerates renal fibrosis. In the presence of pre-existing kidney disease, fructose intake further worsens renal function, increases proteinuria, and enhances glomerulosclerosis. Uric acid has been proposed as a key mediator linking fructose metabolism to oxidative stress, inflammation, and renal injury^8^.

Visceral adipose tissue plays a central role in obesity-related kidney damage by acting as an active endocrine organ. In addition to releasing free fatty acids, adipose tissue secretes adipokines and pro-inflammatory cytokines, including interleukins IL-1, IL-6, and IL-8, as well as tumor necrosis factor-α (TNF-α). These mediators, together with macrophage infiltration into adipose tissue, sustain a chronic low-grade inflammatory state that contributes to renal endothelial dysfunction and progressive parenchymal injury^9^.

The renin–angiotensin–aldosterone system (RAAS) is a critical regulator of renal inflammation and fibrosis in obesity and metabolic disease^10^. RAAS activation modulates intracellular cytokine signaling and directly regulates the expression of transforming growth factor-β1 (TGF-β1), a central profibrotic cytokine that drives extracellular matrix accumulation through increased type IV collagen synthesis^11^.

Oxidative stress represents a key pathogenic mechanism linking metabolic dysregulation to kidney injury. Hyperglycemia and altered lipid metabolism increase superoxide anion generation, leading to excessive reactive oxygen species (ROS) production. This process promotes vascular dysfunction, accelerates renal aging, and amplifies inflammatory signaling^12^. In the renal interstitium, ROS and pro-inflammatory cytokines enhance TGF-β1 signaling, inducing fibroblast activation and epithelial–mesenchymal transition (EMT) of tubular epithelial cells, ultimately resulting in irreversible renal fibrosis^13^.

There is growing evidence in literature that endogenous sex hormones influence cardiovascular–kidney–metabolic (CKM) syndrome. Moreover, premenopausal and postmenopausal women exhibit distinct patterns of CKM conditions compared with men^14^. Menopause is associated with metabolic changes, including increased visceral adiposity, insulin resistance, and dyslipidemia, which collectively exacerbate the risk of MS and CKD progression^15^. Although hormone replacement therapy can mitigate some metabolic consequences of estrogen deficiency^16^, concerns regarding long-term safety limit its widespread use, underscoring the need for alternative therapeutic strategies. In this context, phytoestrogens derived from soy have emerged as potential modulators of metabolic and inflammatory pathways. In ovariectomized rats exposed to a high-fat diet supplemented with fructose, soy phytoestrogens may represent a promising intervention to attenuate obesity-related metabolic disturbances, suppress inflammatory signaling, and limit RAAS- and TGF-β–mediated renal injury^17^.

According to this context, we hypothesize that dietary supplementation with soy phytoestrogens mitigates obesity-associated kidney injury in ovariectomized rats subjected to a high-fat, high-fructose diet by attenuating the mechanisms implicated in CKD progression.

## Methodology

The experimental protocol in vivo was approved by the ethics committee of the Universidade Federal de São Paulo and was performed in accordance with the Brazilian guidelines for scientific animal care and use^18,19^. Female Wistar rats weighing between 190 and 210 g at 6 weeks of age were randomly assigned to the control group (CTL), the high-fat/high-fructose diet group (DH), and the high-fat/high-fructose diet and isoflavones group (DH+ISO) (Figure 1). The animals were subjected to the experimental protocol for a total of 4 months, which was repeated 3 times at different moments with N = 5 for each group. Chow and water were supplied ad libitum. Food consumption and body weight were monitored weekly to carefully characterize weight gain and calorie intake.

**Figure 1.**
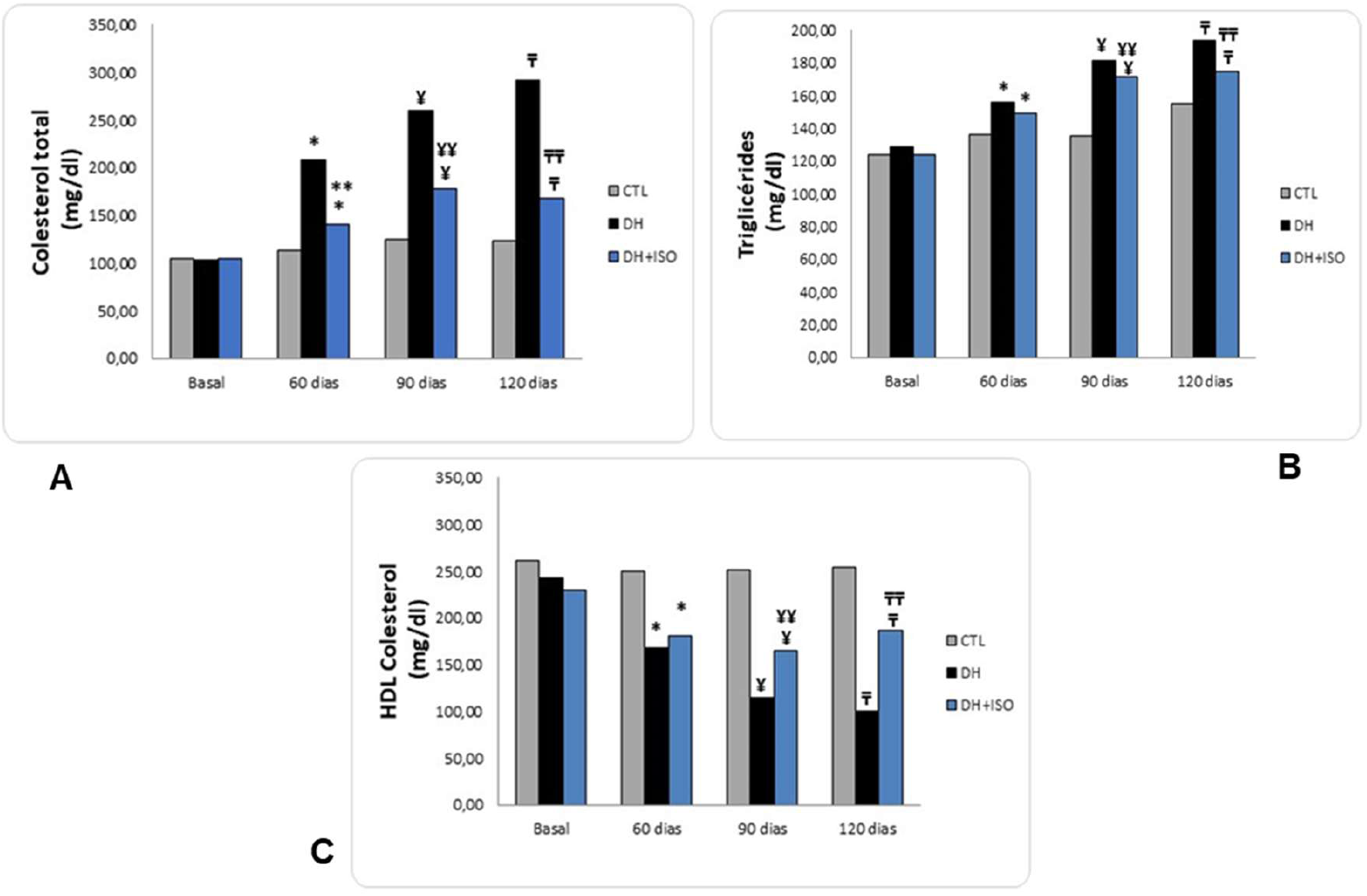
Graphical representation of total cholesterol (A), triglycerides (B), and HDL cholesterol (C) assessment at baseline, 30, 60, 90, and 120 days. Data are reported as mean ± standard error. The significance level for a null hypothesis was set at 5% (p < 0.05). (*) compared to the CTL group at 60 days, (**) compared to the DH group at 60 days, (¥) compared to the CTL group at 90 days, (¥¥) compared to the DH group at 90 days, (τ) compared to the CTL group at 120 days, and (π) compared to the DH group at 120 days.

The animals underwent bilateral ovariectomy surgery at 10-11 weeks of age. The rats were anaesthetized with a solution of ketamine (50mg/kg)/ xylazine (5mg/kg) and isofluorane 4% (respectively) and subsequently submitted to trichotomy of the left paralumbar region (∼ 2.0 cm). The ovary and part of the gonadal adipose tissue were exteriorized and tied with a suture, followed by removal of the ovary. The incision site was sutured and sterilized with iodized alcohol. The same procedure was carried out on the contralateral ovary. At the end of the surgery, the animals were given antibiotics (Pentabiotic, Fort Dodge, dose: 0.2 mL; via intramuscular) and analgesics (Banamine, Schering-Plough; dose: 1mg/kg; via subcutaneous) for adequate recovery from surgery.

The rats were kept under controlled environmental conditions (12/12 h light/dark cycles, and 22–24 ° C), in individual boxes with wood shavings. At baseline, 30, 60, 90, and 120 days after the initiation of the experimental protocol, the animals were placed in metabolic cages for 24 h for urine collection, and blood samples were collected from the lateral tail vein. The rats were euthanized 120 days after the beginning of the experimental protocol through the intraperitoneal injection of a toxic dose of xylazine (10mg/kg)/ketamine (90 mg/kg; Agribands do Brazil Ltda., SP, Brazil), and both kidneys were then removed for immunohistochemistry.

The diets for the respective groups were as follow: the control diet group (CTL) had 20.56% protein, 61.74% carbohydrates, 17.70% lipids, 10% sucrose (glucose and fructose), and 3.9 kcal/g (AIN-93, Rhoster Indústria e Comércio Ltda., SP, Brazil); the high-fat/high-fructose diet group (DH) had 15.32% protein, 16.41% carbohydrates, 68.27% lipids, 20% fructose, 35% lard, and 5.6 kcal/g (RH19543, Rhoster Indústria e Comércio Ltda., SP, Brazil); and the high-fat/high-fructose diet and isoflavones group (DH+ISO) had isoflavones (Ultra Soy Extract, Life Extension, FL, USA) concomitantly administered by gavage with the DH diet during the last 60 experimental days The treatment of animals with ISO was standardized at 300 mg/kg and diluted in propylene glycol, where 45% was isoflavones containing genistein (51.85%), daidzein (40.74%), and glycitein (7.41%).

Metabolic parameters: the rats were assessed by measuring their weight and blood pressure. Body weight was assessed monthly using a scale with a capacity of 2610g and a precision of 1g (Labortex, SP, Brazil) and the result was expressed in g. Systolic blood pressure was measured indirectly by tail plethysmography in awake animals, which were placed in a heated room for 10 minutes, with the cuff and pulse receiver attached to the tail. An electric sphygmomanometer coupled to a two-channel polygraph (Record 2200S, Gould - USA) was used. The results were expressed as an arithmetic mean.

Biochemical Analysis: tests were performed for blood glucose, glycated hemoglobin, total cholesterol and fractions, triglycerides, uric acid, creatinine, creatinine clearance, urea, sodium excretion fraction, and proteinuria, according to the protocol of the commercially available kits (LabTest Diagnóstica) or standardized by the laboratory. The biological material samples were collected every 30 days and stored properly for experimental analysis according to the manufacturer’s protocol. The colorimetric endpoint method consisted of determining the concentration of each analyte through a colorimetric chemical reaction. The reaction was based on the detection of the color developed by the addition of the reagents and the biological sample after incubation. A biochemical analyzer was used, which at the wavelength in visible light (nm) incident light, detecting the absorbance (absorbed light) of each reaction trio: “blank” (standard color, “zero”), standard sample, and test sample. The absorbance was proportional to the analyte concentration, and the results were obtained through the following the calculations recommended by the manufacturer. The results are expressed as percentages.

Analysis of urinary peroxides using the FOX-2 method: measurements of urinary peroxides (FOX-2) were carried out using the xylenol orange version-2 (FOX-2) method, which consists of determining peroxide levels using the iron-xylenol orange method. Peroxides oxidize the Fe2+ ion to the Fe3+ ion when diluted in acidic solutions. Xylol orange [o-cresolsulfonaphthaline 3’, 3”-bis (methylamino) diacetic acid] shows high selectivity for the Fe3+ ion, producing a purplish-blue complex. The FOX-2 solution was prepared using 90 ml (milliliter) of methanol; 10 ml of double-distilled water; 100 µM (micromolar) xylenol orange; 4 mM BHT (2[6]-di-tert-butyl-p-cresol); 25 mM (millimolar) of sulfuric acid solution and 250 µM of ammonium ferrous sulfate. Next, 100 µl (microliter) of urine was added to 900 µl of the FOX-2 solution. The solution was homogenized, kept at room temperature for 30 minutes and centrifuged to remove protein residues. Readings were taken by spectrophotometry at an absorbance of 560 nm (nanometer). The values were stabilized per gram of urinary creatinine and expressed as nmol of peroxides/gram of urinary creatinine.

Histochemical stains on kidney tissue slides were carried out at the Histopathology Laboratory in the Pathological Anatomy Center of the Pathology Center of the Adolfo Lutz Institute (São Paulo, Brazil), by the technician in charge of the sector and blindly. Routine Hematoxylin-Eosin staining was carried out for the histological evaluation, after the histological processing of the kidneys: embedding, cutting histological, deparaffinization and hydration of the tissues. The slides were stained for 2 minutes in Harris hematoxylin and 2 minutes in eosin. After staining, the slides were dehydrated in alcohol and xylene baths, mounted and viewed under optical microscope. In order to assess the deposition of glomerular and tubulointerstitial collagen, Masson’s Trichrome staining was carried out, where the main aspects were: 1 hour in Bouin’s liquid; ferric hematoxylin for 10 minutes; Brebich scarlet for 5 minutes; phosphotungstic acid phosphomolybdic acid for 10 minutes; light green for 5 minutes; glacial acetic acid 1% 3-5 minutes. After staining, the slides were dehydrated in alcohol and xylene baths, mounted and viewed under an optical microscope.

A total of 100 areas were analyzed along the entire kidney section per animal, in an attempt to establish a semi-quantitative evaluation for the following parameters: 1-Acute tubular necrosis (necrosis of renal tubular epithelial cells, with loss of nucleus, granulation or rarefaction of the cytoplasm): 0-3; 2-Tubular dilatation (dilatation of the lumens of the renal tubules in the cortical region, different from the usual histological pattern in rodents, in which the lumen is almost imperceptible under light microscopy): 0-3; 3-Intratubular cylinders (amorphous hyaline/eosinophilic material, compact or granular, inside the renal tubules): 0-3; 4-Interstitial inflammatory infiltrate (presence of inflammatory cells mononucleated in the renal interstitium, perivascular and peritubular): 0-3; 5-Tissue necrosis (altered cell contours, fragmentation of nuclear DNA, with loss of dye affinity to hematoxylin, with tissue eosinophilia): present or absent; 6-Pyelonephritis (presence of inflammatory cells, including polymorphonuclear cells inside the renal tubules, with aggression to the renal epithelium and with extravasation of inflammatory cells into the peritubular interstitium): present or absent. 7-Chronic kidney disease (presence of glomerular hyalinization, tubular atrophy, interstitial tubule fibrosis and lymphomononuclear inflammatory infiltrate): present or absent.

The immunohistochemical examination was carried out in the Immunohistochemistry Laboratory of the Pathological Anatomy Center at the Pathology Center of the Adolfo Lutz Institute. The kidney tissue slides were deparaffinized in xylene baths and rehydrated in gradient alcohol. Antigen recovery was carried out using moist heat in the conventional pressure cooker for 3 minutes with 10mM citric acid pH 6.0. The endogenous peroxidase activity was blocked with 10% H_2_O_2_ for 30 minutes at 37°C. The slides were incubated in a humid chamber with the primary antibodies, described in Table 3, for 18 hours at 4°C. The secondary antibody was then applied for 40 minutes at 37°C and amplified with peroxidase-conjugated polymer for 40 minutes at 37°C (REVEAL Polyvalent HRP detection system, Spring Bioscience). Marking was detected by exposing the slides to the chromogenic substrate DAB (3-3’ diaminobenzidine), and where antigen-antibody binding occurred, a brownish precipitate was generated. The slides were counterstained with Harris hematoxylin, dehydrated in alcohol and xylene baths, mounted and viewed under an optical microscope. The following antibodies were used in the immunohistochemical: CD68 - Dako, Code M 0814 - Denmark), IL-6 (anti-IL-6 - MilliporeMerck, Code CBL2117 - Germany), IL-1β - anti-IL-1β - Santa Cruz, Code sc-7884 (Europe) in kidney tissue samples from ovariectomized rats.

For the statistical analysis, analysis of variance (ANOVA - two way) was used to determine whether there was a significant difference between the means of the three sets of data, followed by Tukey’s post-hoc test to locate the differences. The means were presented as standard error of the means (SEM). A significance level of 0.05 was used for the variables analyzed (p< 0.05).

## Results

### 3.1 Metabolic parameters

Body weight (Table 1) did not differ between DH and DH+ISO groups up to 90 days. At 120 days, body weight increased in the DH group compared with earlier time points, whereas no change was observed in the DH+ISO group. At 120 days, both DH and DH+ISO groups exhibited higher body weight compared with the CTL group.

**Table 1.**
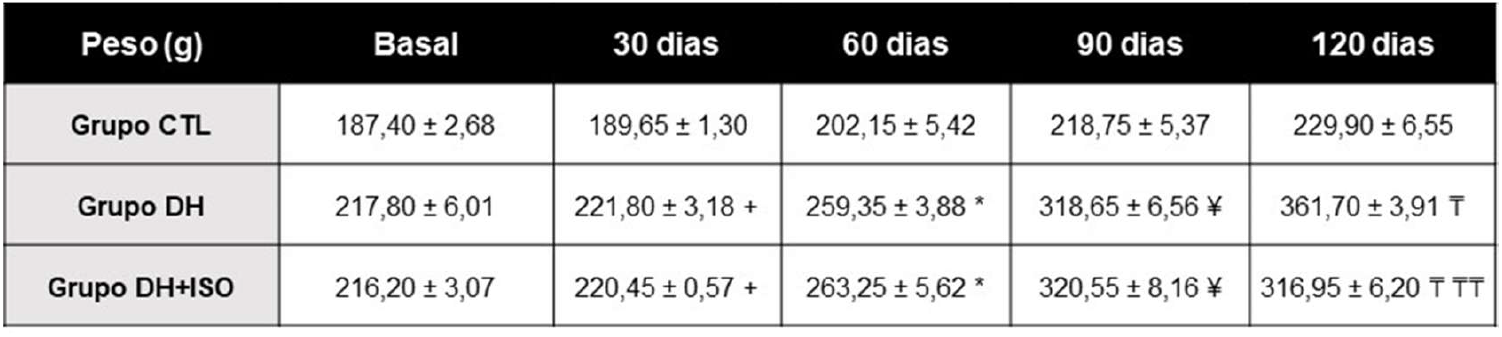
Weight (g) in rats in the control group (CTL), high-fat/high-fructose diet group (DH), and high-fat/high-fructose diet and isoflavones group (DH+ISO) at baseline, 30, 60, 90, and 120 days. Data are reported as mean ± standard error. The significance level for a null hypothesis was set at 5% (p < 0.05). (+) compared to the CTL group at 30 days, (++) compared to the DH group at 30 days, (*) compared to the CTL group at 60 days, (**) compared to the DH group at 60 days, (¥) compared to the CTL group at 90 days, (¥¥) compared to the DH group at 90 days, (τ) compared to the CTL group at 120 days, and (π) compared to the DH group at 120 days.

Systolic blood pressure (SBP) (Table 2) was higher in both DH and DH+ISO groups compared with the CTL group from the third week onward. SBP did not differ between DH and DH+ISO groups until the ninth week. After this time point, SBP decreased in the DH+ISO group, whereas elevated levels persisted in the DH group.

**Table 2.** Systolic blood pressure and systemic hemodynamics. Rats in the CTL group, the DH group, and the DH+ISO group. (A) Measurement of the systolic blood pressure (SBP) in rats at baseline, 3, 5, 7, 9, 11, 13, and 15 weeks. The arrow shows a significant decrease in SBP in the DH+ISO group when compared with DH group at 13 and 15 weeks. Data are reported as mean ± standard error. The significance level for a null hypothesis was set at 5% (p < 0.05). (+) compared to the CTL group at 30 days, (++) compared to the DH group at 30 days, (*) compared to the CTL group at 60 days, (**) compared to the DH group at 60 days, (¥) compared to the CTL group at 90 days, (¥¥) compared to the DH group at 90 days, (τ) compared to the CTL group at 120 days, and (π) compared to the DH group at 120 days.

| PAS (mmHg) | Grupo CTL | Grupo DH | Grupo DH+ISO |
| --- | --- | --- | --- |
| Basal | 144,60 ± 2,34 | 142,80 ± 2,58 | 141,40 ± 2,71 |
| 3 semanas | 145,20 ± 1,43 | 143,60 ± 3,60 | 139,80 ± 1,66 * |
| 5 semanas | 143,60 ± 2,69 | 158,40 ± 3,23 * | 157,80 ± 3,43 * |
| 7 semanas | 143,60 ± 1,60 | 167,80 ± 2,97 * | 166,20 ± 2,56 * |
| 9 semanas | 143,80 ± 3,67 | 165,00 ± 2,35 * | 167,00 ± 2,30 * |
| 11 semanas | 142,80 ± 1,20 | 163,60 ± 1,89 * | 159,60 ± 6,68 * |
| 13 semanas | 142,00 ± 1,87 | 171,20 ± 2,91 * | 148,80 ± 0,86 * ¥ |
| 15 semanas | 144,40 ± 2,91 | 173,20 ± 1,50 * | 153,80 ± 1,98 * ¥ |

### 3.2 Biochemical analysis

Total cholesterol levels (Figure 1A) were higher in the DH and DH+ISO groups compared with the CTL group from 60 days onward. At 90 and 120 days, total cholesterol was higher in the DH group compared with the DH+ISO group. Despite this significant reduction, cholesterol levels in the DH+ISO group remained higher than in the CTL group at all evaluated time points.

Triglyceride levels (Figure 1B) were increased in both DH and DH+ISO groups compared with the CTL group from 60 days onward. At 90 and 120 days, triglyceride levels were higher in the DH group compared with the DH+ISO group. At 120 days, triglyceride levels in the DH+ISO group remained higher than in the CTL group.

HDL cholesterol levels (Figure 1C) were lower in both DH and DH+ISO groups compared with the CTL group from 60 days onward. At 120 days, HDL levels in the DH+ISO group were higher than those observed in the DH group but remained lower than in the CTL group.

Plasma glucose levels (Figure 2A) were higher in both DH and DH+ISO groups compared with the CTL group between 30 and 120 days. From 90 days onward, glucose levels were lower in the DH+ISO group compared with the DH group.

**Figure 2.**
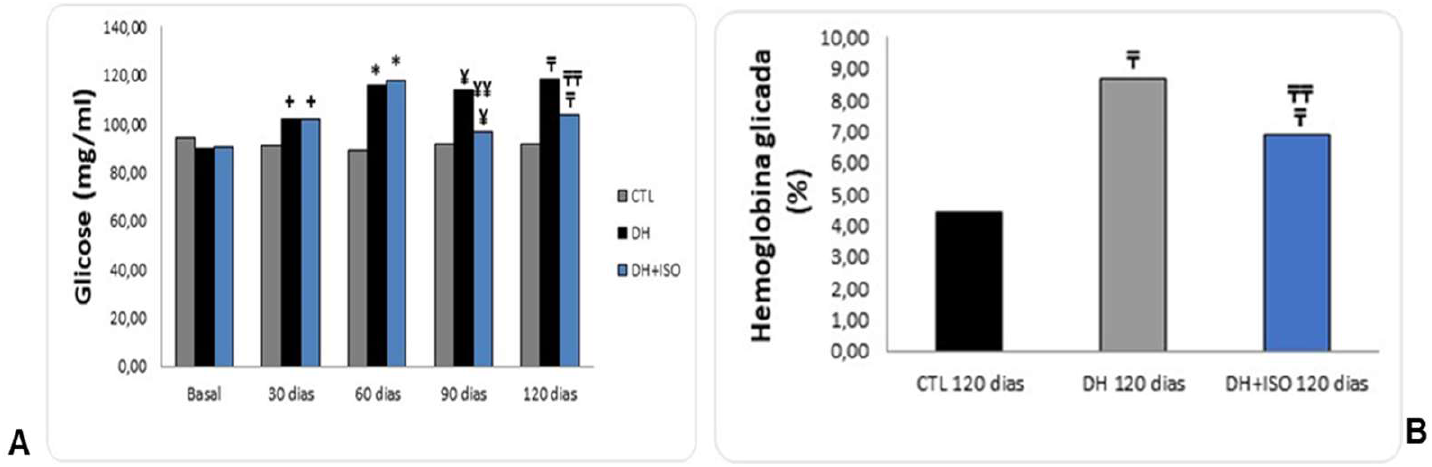
Graphical representation of glucose levels (A) and Glycated hemoglobin (B), assessment at baseline, 30, 60, 90, and 120 days. Data are reported as mean ± standard error. The significance level for a null hypothesis was set at 5% (p < 0.05). (*) compared to the CTL group at 60 days, (**) compared to the DH group at 60 days, (¥) compared to the CTL group at 90 days, (¥¥) compared to the DH group at 90 days, (τ) compared to the CTL group at 120 days, and (π) compared to the DH group at 120 days.

Glycated hemoglobin (Figure 2B), evaluated at 120 days, was higher in the DH group compared with both the CTL and DH+ISO groups. HbA1c levels in the DH+ISO group were higher than in the CTL group and did not differ from those observed in the DH group.

Plasma creatinine levels (Figure 3A) were higher in both DH and DH+ISO groups compared with the CTL group at 90 and 120 days. At 120 days, plasma creatinine levels were lower in the DH+ISO group compared with the DH group.

**Figure 3.**
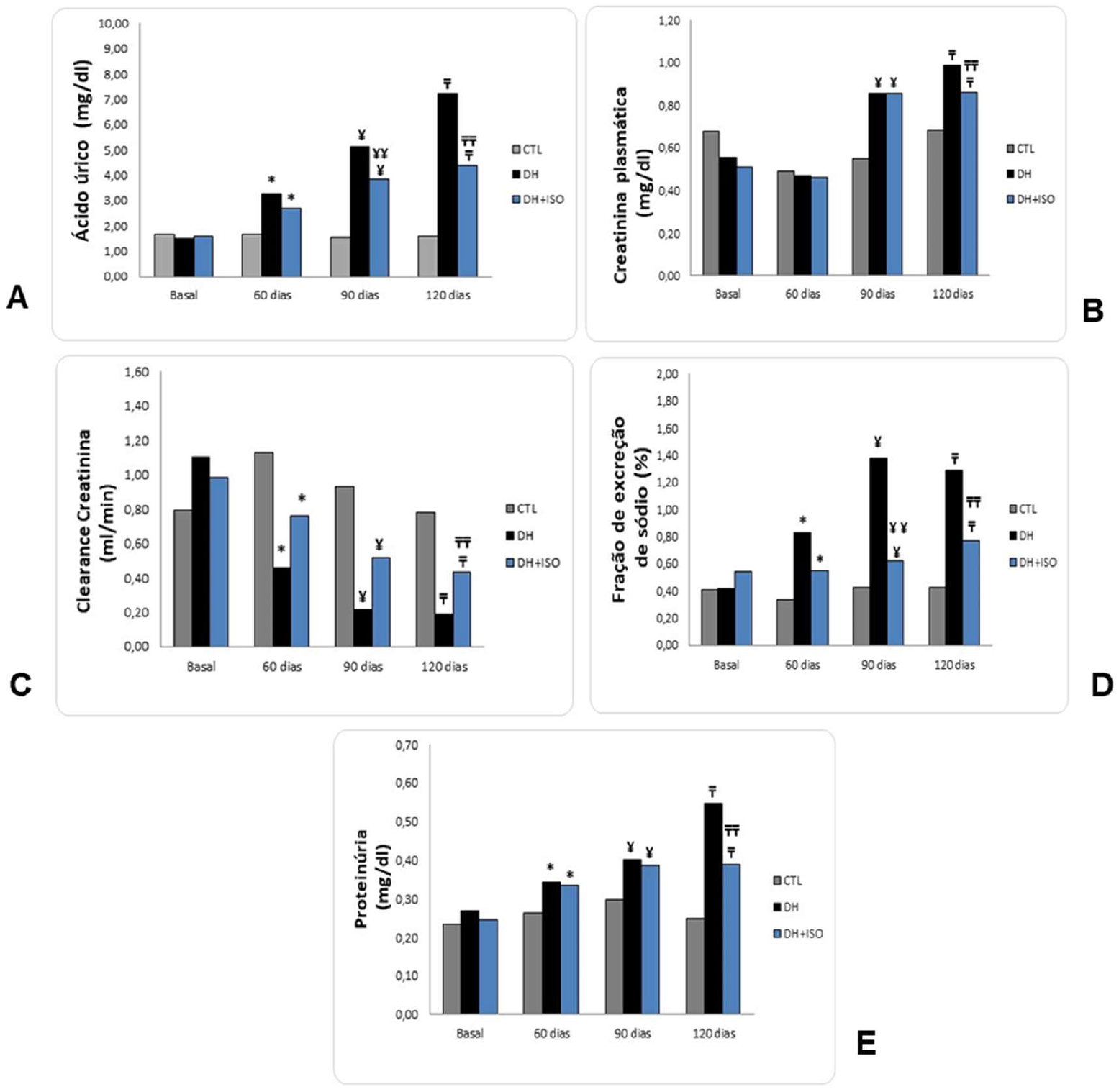
Graphical representation of plasma creatinine (A), creatinine clearance (B), urea (C), sodium excretion fraction (D) and proteinuria (E) assessment at baseline, 30, 60, 90, and 120 days. Data are reported as mean ± standard error. The significance level for a null hypothesis was set at 5% (p < 0.05). (*) compared to the CTL group at 60 days, (**) compared to the DH group at 60 days, (¥) compared to the CTL group at 90 days, (¥¥) compared to the DH group at 90 days, (τ) compared to the CTL group at 120 days, and (π) compared to the DH group at 120 days.

Creatinine clearance (Figure 3B) was lower in the DH group compared with the CTL group at all time points. Creatinine clearance was higher in the DH+ISO group compared with the DH group at 90 and 120 days but remained lower than in the CTL group.

Plasma urea levels (Figure 3C) were higher in the DH+ISO group compared with the CTL group at 90 days and higher in the DH group compared with the CTL group at 120 days. At 120 days, urea levels in the DH+ISO group did not differ from those in the CTL group.

The fractional excretion of sodium (Figure 3D) was higher in the DH group compared with the CTL group from 60 days onward. Values in the DH+ISO group were higher than in the CTL group and did not differ from those observed in the DH group.

Proteinuria (Figure 3E) was higher in both DH and DH+ISO groups compared with the CTL group at 120 days. At earlier time points (60 and 90 days), proteinuria was higher in the DH+ISO group compared with the CTL group.

Urinary peroxide levels (FOX-2) (Figure 4) were higher in both DH and DH+ISO groups compared with the CTL group from 60 to 120 days. At 120 days, FOX-2 levels were lower in the DH+ISO group compared with the DH group.

**Figure 4.**
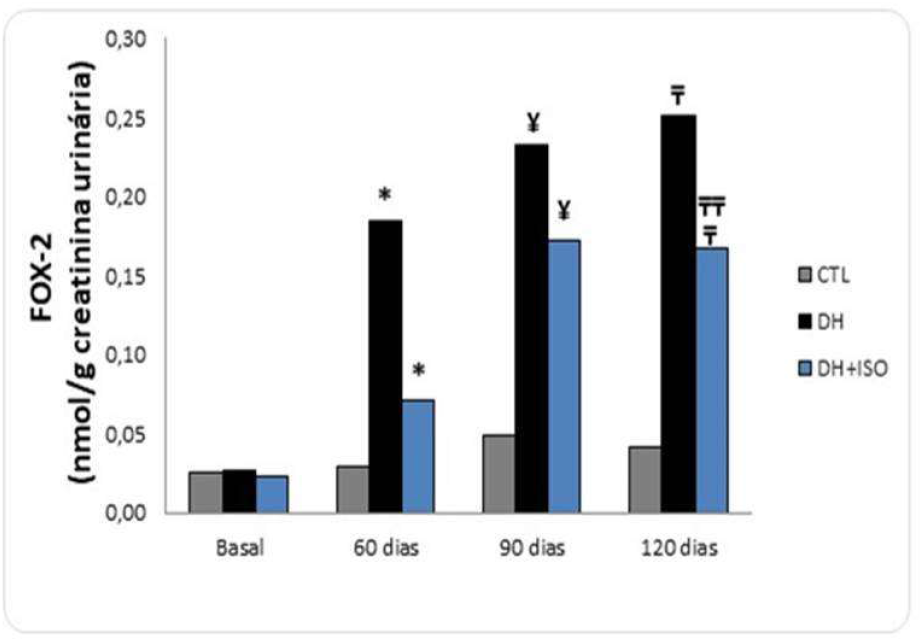
Graphical representation of the assessment of urinary peroxide levels (FOX-2) assessment at baseline, 30, 60, 90, and 120 days. Data are reported as mean ± standard error. The significance level for a null hypothesis was set at 5% (p < 0.05). (*) compared to the CTL group at 60 days, (**) compared to the DH group at 60 days, (¥) compared to the CTL group at 90 days, (¥¥) compared to the DH group at 90 days, (τ) compared to the CTL group at 120 days, and (π) compared to the DH group at 120 days.

### 3.3 Histopathological analysis

The histological analysis of the experimental groups (CTL, DH, and DH+ISO) was represented by semiquantitative analyses, establishing scores for relevant findings in the analysis of renal morphology (0 = absent; 1 = mild, 2 = moderate, and 3 = severe) (Figue 5A-D). In the CTL group, we detected slight tubular dilatation (score 1) in 100% (5/5) of the animals, acute tubular necrosis in 100% (5/5), slight cylinder formation in 60% (3/5), and 1 of the animals (1/5) in the CTL group presenting with severe renal failure (score 3), detected by the presence of pyelonephritis. This animal is an outlier, and the presence of pyelonephritis is not related to the diet and was not observed in the other animals, where we did not detect any noteworthy histopathological changes (Figure 5A).

**Figure 5.**
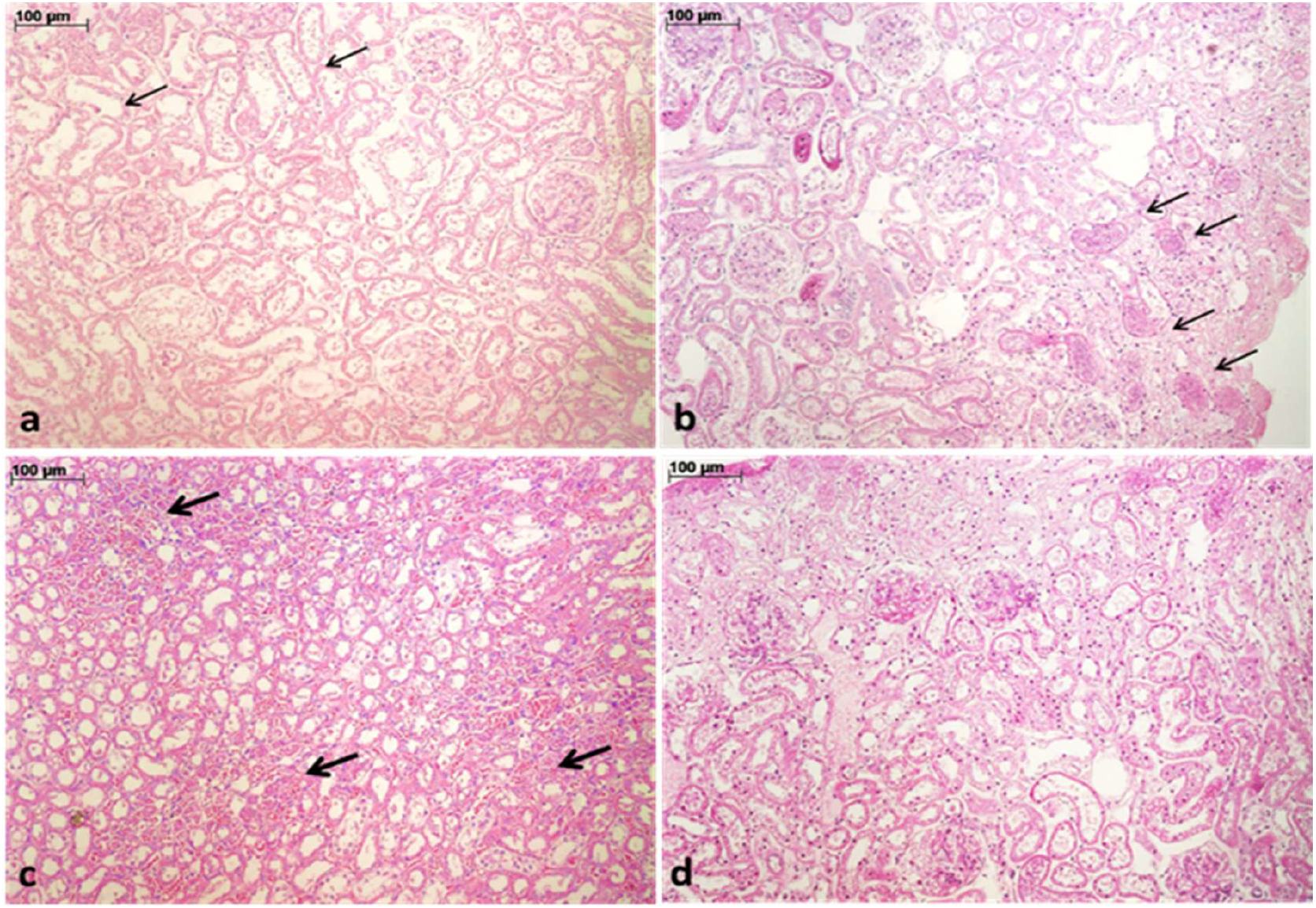
Histological findings in rats after ovariectomy and subjected to a high-fat diet, with or without isoflavone, and control animals. A-renal cortex of control rats on a standard diet, showing mild acute tubular necrosis (tubular dilation and loss of nucleus in some epithelial cells); B and C-renal cortex of rats subjected to a high-calorie diet supplemented with fructose, showing intense tubular necrosis and numerous granular casts in the tubular lumen (b, arrows) and foci of necrosis in distal tubules, near the renal pelvis (c, arrows); D-cortex of a rat subjected to a high-calorie diet supplemented with fructose and treated with isoflavone, showing acute tubular necrosis, moderate with rare tubular casts. Staining: Hematoxylin-Eosin. Magnification: 100x, A-D. Scale bar: 100μm, A-D.

The DH group showed moderate tubular dilatation (score 2) in 100% (5/5) of the animals, moderate acute tubular necrosis in 80% of the animals (4/5), and 1 animal (1/5) with severe acute tubular necrosis (score 3), moderate in 100% (5/5) of the animals, no evidence of renal failure, and the presence of cortical necrosis in 80% (4/5) of the animals as the main histopathological change (Figure 5B-C).

The DH+ISO group showed moderate tubular dilatation (score 2) in 100% (5/5) of the animals, moderate acute tubular necrosis (score 2) in 80% of the animals (4/5), and 1 animal (1/5) with mild acute tubular necrosis (score 1), mild cylinder formation (score 1) in 40% (2/5) of the animals and 60% (3/5) with no cylinders (granular), and the presence of cortical necrosis in 80% (4/5) of the animals as the main histopathological change (Figure 5D).

The DH and DH+ISO groups exhibited higher scores for tubular dilatation, acute tubular necrosis, and cortical necrosis compared with the CTL group. No difference was observed between DH and DH+ISO groups (Figure 5).

### 3.4. Immunohistochemical examination

The evaluation of IL-1β by immunohistochemical staining showed a decrease in the expression of the proinflammatory cytokine in the DH+ISO group compared to the group without isoflavone administration (DH group) (Figure 6A), but there was no statistically significant difference between the DH and DH+ISO groups (p>0.05).

**Figure 6.**
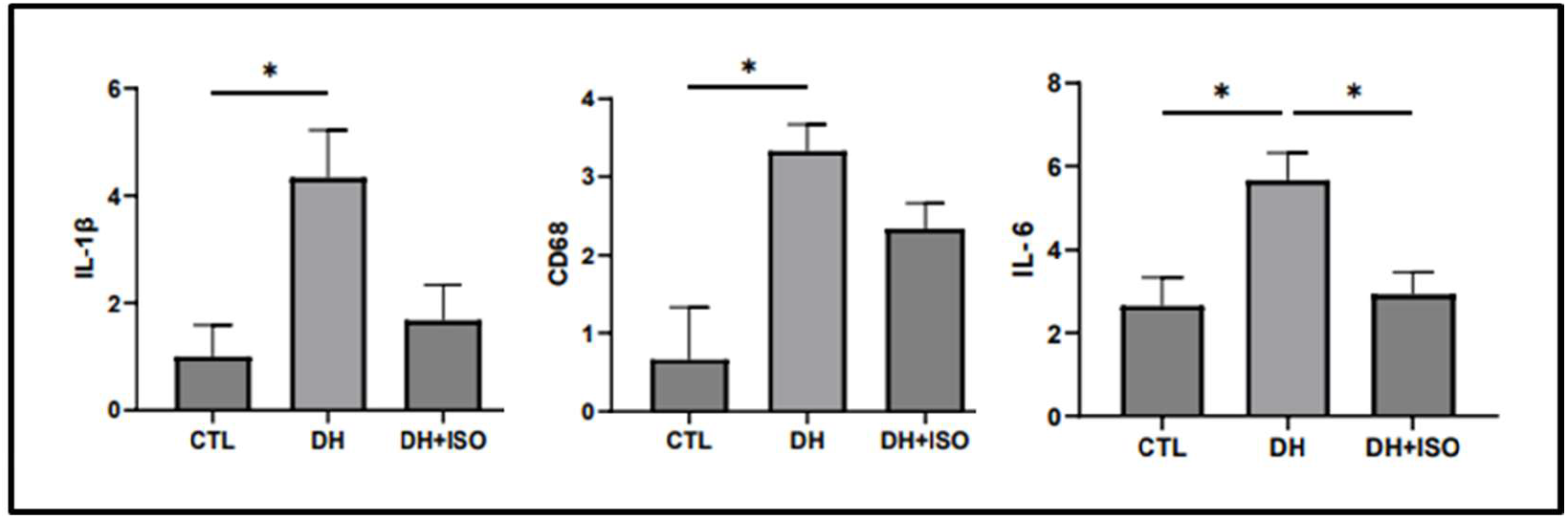
Graph representing the detection of IL-1β, CD68, and IL-6 through immunohistochemical staining in renal tissue, obtained by reading 20 microscopic fields with a final magnification of 400x. Results area expressed as cell numbers. The comparison between the control (CTL), untreated (DH), and treated (DH+ISO) groups showed a statistically significant difference between the CTL and DH groups, and there was a statistically significant difference between the DH and DH+ISO groups (p<0.05). 624

Macrophage and monocyte infiltration was assessed by counting the number of CD68-positive cells in the kidney. Immunohistochemical staining for CD68 showed a decrease in macrophages in the DH+ISO-treated group compared to the group without isoflavone administration (DH group) (Figure 6B). There was no statistically significant difference between the DH and DH+ISO groups (p>0.05).

The evaluation of IL-6 by immunohistochemical staining showed a decrease in the expression of proinflammatory cytokine in the DH+ISO treated group (Group 3) compared to the group without isoflavone administration (Group 2) (Figure 6C), with a statistically significant difference between the groups (p<0.05). 311

## 4. Discussion

Modern diets are characterized by ultra-processed foods that are high in sugar, fat, and calories. In addition, sedentary lifestyles contribute to rising obesity rates. This chronic medical condition is not limited to excess weight; it is also associated with cardiovascular disease, type II diabetes, musculoskeletal disorders, and even mental health problems such as depression and anxiety^20,21^.

According to the 2023 World Obesity Atlas, by 2035 obesity and overweight are expected to affect half of the world’s adolescents and children, equivalent to two in five children globally. According to this document, most people who live and die with chronic noncommunicable diseases (NCDs) are overweight or obese^22^.

In middle-aged women, during the transition to menopause, this is common, and the adverse effects resulting from these changes are a point of discussion in clinical practice, since hormonal changes result in a marked gain in central body fat^23^. This fact may be exacerbated by the Western diet and lack of physical activity.

In recent years, fructose has been used as a sweetener in sweet drinks and corn syrups, justified by its greater sweetening power when compared to glucose and because it is more palatable. However, its increased use in industrialized products and indiscriminate consumption may trigger various metabolic disorders^24^.

The combination of a high-fat diet and fructose consumption further increased weight gain when compared to a high-fat, high-glucose diet, resulting in resistance to the hormone leptin and accelerating obesity, i.e., a condition of leptin resistance predisposing to the development of dietary obesity^25^.

Our experimental model in ovariectomized rats served to mimic what happens after menopause, with the drop in natural estrogen, a new distribution of body fat occurs, with the accumulation of visceral fat, insulin resistance, and dyslipidemia, two important factors for the characterization of MS, in addition to renal dysfunction and cardiovascular problems.

The impact of a high-fat diet supplemented with fructose was an important factor in the development of obesity and MS in the animals in our study, with significant changes in some of the metabolic and biochemical parameters evaluated. This may reflect what happens in this population, which follows a modern diet rich in fat and sugars in their daily lives^3^.

The animal models described in the literature and subjected to DH developed MS with changes in some important biochemical markers such as triglycerides, insulin, and uric acid^26^. Thus, in our experimental model, rats subjected to the DH diet without isoflavone administration developed MS, as evidenced by changes in metabolic and biochemical markers such as weight, lipid profile, blood glucose, and impaired renal function.

One reason why kidneys are affected by diets high in fructose is related to its catabolism. Fructolysis can lead to depletion of intracellular energy, mitochondrial oxidative stress, and production of inflammatory mediators. Uric acid plays this role, so hyperuricemia is related to impaired kidney function^27^.

In a study published by Rebholz et al. in 2019, in a cohort of African Americans, it was found that high consumption of sugary drinks results in a higher probability of developing CKD^28^. In our study, some of the histopathological changes detected may progress to CKD.

In our experimental model, the renal histopathological changes found in the DH diet group resulted in cortical necrosis and the detection of granular casts, findings that are related to acute kidney disease. It should be noted that one of the renal pathological findings described in the literature in animals subjected to DH is related to tubulointerstitial injury, the latter histopathological finding was observed in our experimental group subjected to DH^**8**^.

The presence of cortical necrosis will likely trigger a tissue repair mechanism, resulting in long-term renal fibrosis and a diagnosis of CKD. Tubular necrosis may have been the result of lipid deposition and its nephrotoxic effect, in addition to the detection in our study of an increase in EROS, stimulated by lipotoxicity in obese animals, since these substances have already been described in the literature as responsible for apoptosis and cell membrane damage^29,30^.

In a study conducted by Aoyama et al. (2012), three strains of mice were fed a diet of equal caloric value, with only one of them receiving added fructose. The histopathological findings after eight weeks already indicated inflammation, renal fibrosis in the outer cortex, and the presence of oxidative stress. This was justified by the expression of GLUT5 to fructose in the kidney and the role of this expression in the progression of kidney disease and the absence of glomerular kidney changes. In our study, glomerular changes were less evident, but changes in the renal cortex were clearly present in the DH group through findings of cortical necrosis^31^.

In this study, the proposed treatment for remedying damage caused by a high-fat diet supplemented with fructose was the use of isoflavone and its benefits in cases of MS and renal function. It has already been described in the literature that the administration of isoflavone can contribute to minimizing obesity/MS and kidney damage, mainly by mitigating the impact of the modern diet, and is promising for postmenopausal and obese women^32,33^.

The isoflavone-supplemented diet was evaluated in the treatment of MS and obesity by Banz et al. (2004), and their results indicated that obese rats showed a reduction in body weight and plasma lipids^34^. The animals in our experimental group, treated with isoflavone, managed to lose weight, in addition to showing improvement in their glycemic and lipid profiles, including an increase in HDL, in rats treated with isoflavone after 90 days on a DH diet, data that corroborate the literature.

In a study published by Choi and Song (2009), the effect of genistein in ovariectomized rats produced an increase in serum estrogen levels and renal expression of alpha and beta receptors, with a decrease in serum insulin, triglyceride, and cholesterol levels. The administration of genistein prevented severe damage to renal function by reducing insulin resistance, renal oxidative stress, and the accumulation of lipids and extracellular matrix proteins. It is worth noting that we detected a decrease in urinary peroxides in our DH+ISO group, evidencing an improvement in renal oxidative stress^35^.

In the renal histopathological evaluation, we observed that a smaller number of animals treated with isoflavone developed cortical necrosis. We did not detect the formation of granular casts in this group, indicating normal renal flow and no tubular cell degradation or inflammatory cells.

We believe that the administration of isoflavone had a renoprotective effect, as there was a reduction in inflammation and renal impairment in the study animals, verified by the decrease in IL-6 in the group treated with isoflavones and by the less severe histopathological changes, given that processes harmful to the kidney are potentiated in the presence of IL-6^36^.

The histological changes found in the animals in this study are precursors of severe renal involvement. Even in this 120-day experimental model, there are already changes in blood pressure and important microvascular changes that are precursors of chronic disease.

In a 2016 study by Jing and Wei-Jie, CKD patients who consumed soy protein showed improvement in laboratory markers such as decreased serum creatinine, serum phosphorus, C-reactive protein, and proteinuria. Although promising, this study suggests the need for further evaluation of patients using soy/isoflavones, as no significant changes were observed in creatinine clearance, renal clearance rate, body weight, and BMI^37^.

It is believed that excessive fructose metabolism, resulting from modern diets, may be related not only to increasing rates of obesity, but also to hypertension, hepatic steatosis, vascular changes, and even aging. Therefore, conducting studies aimed at remedying this situation becomes extremely relevant^38^.

In a study published by Yanai et al. (2021), hyperuricemia is associated with worsening MS symptoms, which can result in CKD and cardiovascular disease. This study shows us the seriousness of the association between a fructose-rich diet and worsening MS symptoms^39^. In our study, we demonstrated the impact of this diet associated with menopause, and our results corroborated the aforementioned study.

Looking ahead, there is a need to delve deeper into the impact of modern diets on different stages of adulthood, especially during menopause, particularly when associated with aggravating conditions.

The evaluation of new therapeutic strategies, such as the use of nutraceuticals, is a promising strategy and could be incorporated as a complementary therapy in public health, thanks to the results obtained in our work, minimizing MS and renal impairment.

In 2023, the American Heart Association emphasized the importance of further studies on metabolic risk factors, aiming at the prevention and treatment of cardiovascular-renal-metabolic syndrome and its impacts on the population, given that there is heterogeneity in clinical phenotypes, interactions between social determinants of health, and biological risk factors that vary widely in global distribution, challenging scientific understanding^1^.

The administration of isoflavone contributed to the reduction of laboratory markers, inflammatory markers, oxidative stress markers, and renal histopathology, and may be a promising therapeutic strategy not only against renal impairment, but also in cardiovascular-renal-metabolic syndrome.

## Author Contributions

Writing - Original Draft Investigation, TSL; Validation, EAP, FTB; Methodology, TSL, EAP, ANDN and CSCB; Investigation, TSL, EAP, BC, ASO, AACS, and CD; Writing-Reviewing and Editing, TSL, MBC and FTB; Formal analysis; Visualization, TSL and FTB; Resources, AC and MFV; Funding acquisition, MFV, FTB; Project administration, FTB; Conceptualization, FTB; Supervision, FTB. All authors have read and agreed to the published version of the manuscript.

## Acknowledgments

Conselho Nacional de Desenvolvimento Científico Tecnológico (CNPq), Universidade Estudos e Projetos (FINEP), Fundação Oswaldo Ramos (FOR), Fundação de Amparo à Pesquisa do Estado de São Paulo (FAPESP), and Coordenação de Aperfeiçoamento de Pessoal de Nível Superior (CAPES).

## Funding

Fundação de Amparo à Pesquisa do Estado de São Paulo (FAPESP 2020/13405-2).

## Conflicts of Interest

The authors declare no conflict of interest.

## Notes

### Competing Interest Statement

The authors have declared no competing interest.

